# Vestibular-derived internal models in active self-motion estimation

**DOI:** 10.1101/2024.04.14.589435

**Authors:** Milou J.L. van Helvert, Luc P.J. Selen, Robert J. van Beers, W. Pieter Medendorp

## Abstract

Self-motion estimation is thought to depend on sensory information as well as on sensory predictions derived from motor feedback. In driving, the vestibular afference can in principle be predicted based on the steering motor commands if an accurate internal model of the steering dynamics is available. Here, we used a closed-loop steering experiment to examine whether participants can build such an internal model of the steering dynamics. Participants steered a motion platform on which they were seated to align their body with a memorized visual target. We varied the gain between the steering wheel angle and the velocity of the motion platform across trials in three different ways: unpredictable (white noise), moderately predictable (random walk), or highly predictable (constant gain). We examined whether participants took the across-trial predictability of the gain into account to control their steering (internal model hypothesis), or whether they simply integrated the vestibular feedback over time to estimate their travelled distance (path integration hypothesis). Results from a trial series regression analysis show that participants took the gain of the previous trial into account more when it followed a random walk across trials than when it varied unpredictably across trials. Furthermore, on interleaved trials with a large jump in the gain, participants made fast corrective responses, irrespective of gain predictability, suggesting they also rely on vestibular feedback. These findings suggest that the brain can construct an internal model of the steering dynamics to predict the vestibular reafference in driving and self-motion estimation.

## Introduction

Sensory feedback, especially visual and vestibular, is important in self-motion estimation. People can estimate their self-motion from visual (Britten, 2008) or vestibular (Cheng & Gu, 2018) information alone, but are more precise when feedback from both is available and integrated (DeAngelis & Angelaki, 2011; ter Horst *et al*., 2015; Medendorp & Selen, 2017; Britton & Arshad, 2019; Keshavarzi *et al*., 2023).

When the motion is generated actively, self-motion estimates also depend on predictions from internal models of sensory and body dynamics that transform motor commands into predicted sensory consequences (Laurens & Angelaki, 2017; Brooks & Cullen, 2019). In combination with actual sensory feedback, these predictions lead to better estimates of self-motion (Medendorp, 2011; Sanders *et al*., 2011; Campos *et al*., 2012; Carriot *et al*., 2013; Genzel *et al*., 2016), also in patients with vestibular deficits (Worchel, 1952; Glasauer *et al*., 2002; Kaski *et al*., 2016; Medendorp *et al*., 2018). Both during passive and active self-motion, the sensory feedback is thought to be continuously monitored in order to update the self-motion estimate and adjust the internal model if necessary (Prsa *et al*., 2015; Brooks *et al*., 2015).

The role of sensory feedback and predictions in self-motion estimation has been studied with closed-loop steering experiments in both monkeys (Roy & Cullen, 2001; Page & Duffy, 2008; Egger & Britten, 2013; Jacob & Duffy, 2015) and humans (Lakshminarasimhan *et al*., 2018; Stavropoulos *et al*., 2022; Alefantis *et al*., 2022; van Helvert *et al*., 2022). In these experiments, the self-motion is controlled by a joystick or steering wheel, and the sensory feedback can in principle be predicted based on the steering motor command if an accurate internal model of the steering dynamics is available. Alefantis et al. (2022) studied human steering behaviour in a virtual environment and found that participants were able to navigate the environment on trials with optic flow cues, but also on interleaved trials without any sensory feedback, suggesting that participants had formed an internal model of the steering dynamics with training. Similarly, Stavropoulos et al. (2022) studied navigation with optic flow and vestibular cues while the steering dynamics varied from trial to trial according to a random walk (i.e., the dynamics on the previous trial are predictive of the dynamics on the current trial), from responsive to sluggish steering control. Their participants could steer accurately whenever optic flow cues were provided, but less so when only vestibular cues were available and steering control was responsive. It is thus not evident that vestibular cues alone can be used to build an internal model by which the generated self-motion can be predicted based on the steering motor command under changing steering dynamics. This is the topic of the present study.

We have previously examined the role of predictions and sensory feedback, in particular vestibular feedback, in self-motion estimation in a steering experiment in which the steering dynamics changed only twice during the experiment (van Helvert *et al*., 2022). Seated on a linear motion sled, participants were instructed to align their body with a memorized visual target using a steering wheel that controlled their lateral body motion. We found that participants responded rapidly (i.e., made within-trial adjustments to their steering movement) to the sudden step changes in the steering dynamics (i.e., the gain between the steering wheel angle and their body velocity). Across trials, participants’ performance gradually improved further by adjusting to the new steering dynamics. One explanation of these findings is that participants built an internal model of the steering dynamics, which transforms the steering motor commands into predicted vestibular feedback, that they continued to update throughout the experiment based on the vestibular feedback (Brooks *et al*., 2015; van Helvert *et al*., 2022). Another explanation is that participants simply relied on path integration mechanisms (Loomis *et al*., 1993; Lappe *et al*., 2007; Zhou & Gu, 2023), estimating their location relative to the target by integrating the vestibular information over time without building an internal model of the steering dynamics. In the present study we aim to distinguish between these two explanations (internal model versus path integration), taking inspiration from studies on the adaptation of reaching movements.

Burge et al. (2008) and Wei and Körding (2010) studied visuomotor adaptation of reaching movements while the uncertainty of the visual feedback about the reach endpoint and the uncertainty of the spatial mapping between the reach endpoint and the visual feedback was varied. It was found that adaptation proceeded slower with higher visual feedback uncertainty and faster with higher spatial mapping uncertainty. Gonzalez Castro et al. (2014) compared adaptation to a force field that varied in strength unpredictably across trials or to a force field that followed a random walk across trials. They found that participants relied more on sensory feedback in the unpredictable condition, while trusting sensory predictions more in the random walk condition.

In the present study, we used a similar experimental design to dissociate the contribution of vestibular feedback and vestibular predictions in self-motion estimation during driving. Participants steered a linear sled on which they were seated to translate their body to a memorized visual target. We varied the gain between the steering wheel angle and the velocity of the sled across trials in three different ways: a white noise condition (unpredictable gain), a random walk condition (moderately predictable gain) and a constant gain condition (highly predictable).

We examined the steering behaviour for within-trial responses to the vestibular feedback and vestibular predictions based on an internal model of the steering dynamics. Furthermore, we assessed the participants’ responses to more extreme changes in the steering dynamics by introducing large jumps in the gain (i.e., step trials) near the end of each trial block. If participants simply integrated the vestibular information over time to estimate the travelled distance (path integration hypothesis), we would expect to see no differences in the steering behaviour across the three conditions. In contrast, if participants did take the across-trial predictability of the gain into account (internal model hypothesis), we expect them to respond fastest to changes in the gain in the white noise condition, followed by the random walk condition and the constant gain condition.

## Methods

### Participants

The study was approved by the ethics committee of the Faculty of Social Sciences of Radboud University Nijmegen, the Netherlands. Twenty-six naïve participants took part in the study (7 men and 19 women; 18-30 years old) and gave their written informed consent before the start of the experiment. They reported to have normal or corrected-to-normal vision, normal hearing, and no history of motion sickness. The experiment took around 90 minutes per participant, and participants were compensated with course credit or €15,00.

### Setup

Participants were seated on a custom-built linear motion platform, also called the sled, and used a steering wheel to control the sled speed (Fig. 1A). They sat with their interaural axis aligned with the motion axis of the sled, such that they were laterally translated. They were restrained by a five-point seat belt and could stop the sled motion at any time by pressing one of the emergency buttons on either side of the sled chair. The experiment was performed in darkness. The sled was powered by a linear motor (TB15N; Tecnotion, Almelo, The Netherlands) and controlled by a servo drive (Kollmorgen S700; Danaher, Washington, DC, USA). The sled track was approximately 93 cm long. The steering wheel (G25 Racing Wheel; Logitech, Lausanne, Switzerland) was mounted at a comfortable handling distance in front of the participant at chest level and had a resolution of 0.0549° and a range of rotation from −450° to +450°. The steering wheel angle was recorded at 100 Hz. Participants viewed a 55-inch OLED screen (55EA8809-ZC; LG, Seoul, South Korea) with a resolution of 1920 x 1080 pixels and a refresh rate of 60 Hz, positioned centrally in front of the sled track at a viewing distance of approximately 170 cm, and wore noise-cancelling earphones to mask auditory cues induced by the sled motion with white noise sounds (QuietComfort 20; Bose Corporation, Framingham, MA, USA). The experiment was controlled using custom-written software in Python (v.3.6.9; Python Software Foundation).

**Figure 1.**
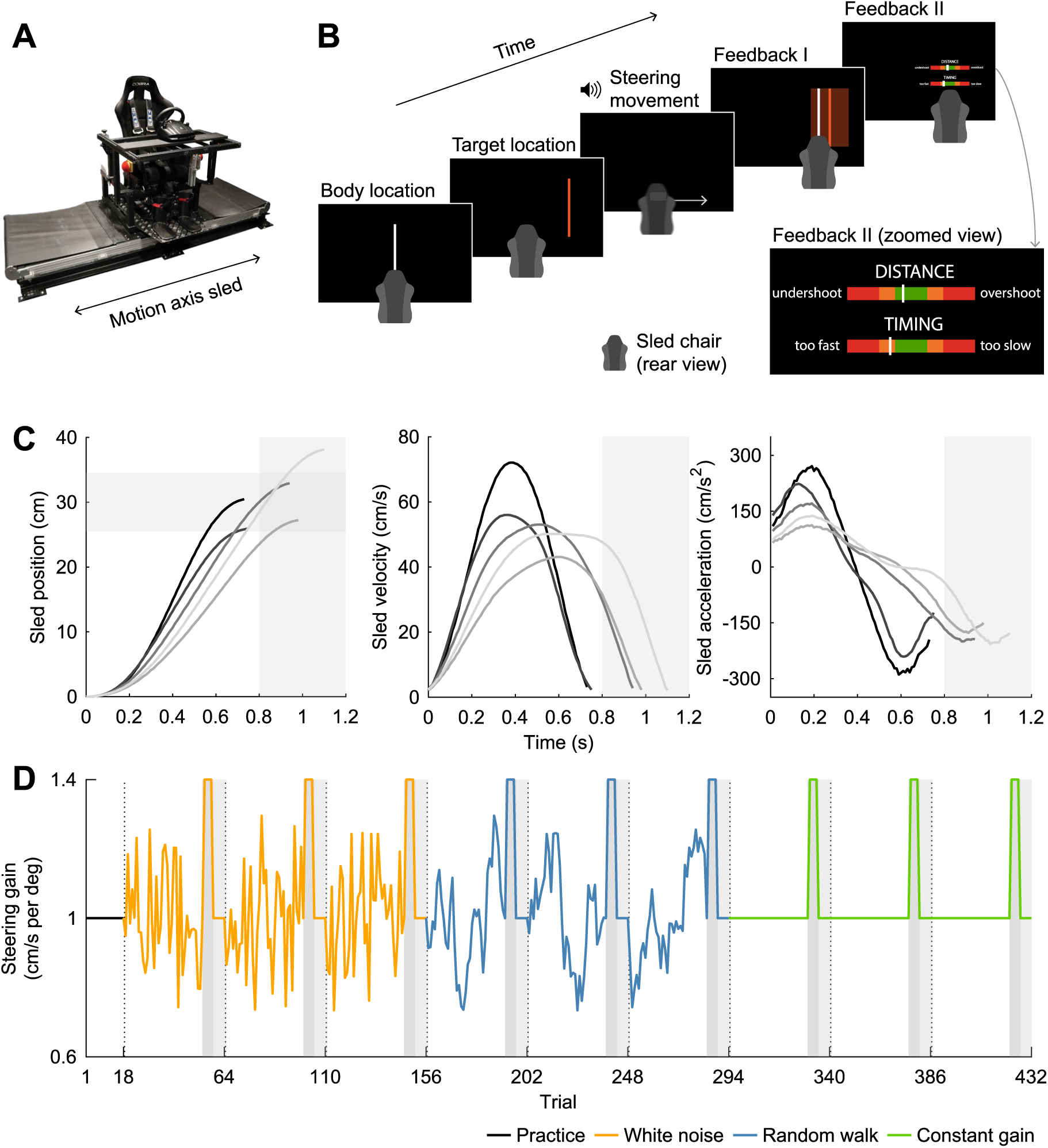
Experimental setup and paradigm. A) Experimental setup. Participants were seated with their interaural axis aligned with the motion axis of the sled and turned a steering wheel to control the sled velocity. B) Experimental paradigm. Participants were first shown their location as a white line, followed by the target location as an orange line. After the disappearance of the target location, a beep instructed participants to turn the steering wheel to translate their body and align it with the memorized target location. When the sled speed was again close to 0 cm/s, visual feedback about the displacement error (Feedback I and II) and the movement duration (Feedback II) was provided. Inset shows the zoomed view of the feedback bars in Feedback II. C) Sled position, velocity and acceleration as a function of time (aligned to movement onset) for five representative condition-specific trials in the constant gain condition (grey lines). For each trial, the measured absolute sled position relative to the start location, the absolute sled velocity encoded by the steering wheel angle, and the sled acceleration, computed by low-pass filtering the derivative of the encoded sled velocity using a moving average filter with a window length of nine samples, are shown. D) Example of the steering gain across trials. Each participant completed nine trial blocks. Each trial block started with 36 condition-specific trials, in which the gain varied from trial to trial (white noise and random walk condition) or remained the same (constant gain condition). Participants were exposed to the exact same gains in the white noise and random walk condition, but trials were organized such that their lag-1 autocorrelation was close to zero in the white noise condition and above 0.8 in the random walk condition. Each trial block was concluded with a baseline trial (gain of 1.0 cm/s per deg), followed by four step trials (high gain of 1.4 cm/s per deg; dark grey areas) and six washout trials (baseline gain of 1.0 cm/s per deg; light grey areas). Participants completed three trial blocks per condition, each followed by a short break (dashed vertical lines), and completed 18 practice trials before the experiment (baseline gain of 1.0 cm/s per deg).

### Paradigm

Figure 1B shows the order of events during an experimental trial. At the start of a trial, a vertical white line aligned with the body midline (width 0.3 cm and height 25.4 cm) was presented on the screen for 1 second, which represented the start location of the body. After this, a vertical orange line (width 0.3 cm and height 25.4 cm) was presented on the screen for 1 second, representing the target location. The target location was alternately presented to the left and to the right of the start location of the body. The target distance, defined as the distance between the start location of the body and the target location, was always 30 cm. Participants were not informed about the fixed target distance.

After disappearance of the target, a short beep was played via the earphones to instruct the participant to rotate the steering wheel to align their body midline with the memorized target location. The sled motion started when the participant turned the steering wheel 0.0549 deg (the smallest detectable change) away from the steering wheel angle at trial start. The steering wheel angle at trial start was typically between −20 and 20 deg, with 0 deg representing the centre of the range. The angle of the steering wheel relative to the angle at trial start encoded the velocity of the sled, but the exact steering gain changed throughout the experiment (see below). The latency between the rotation of the steering wheel and the translation of the sled was approximately 25 ms. The maximum speed of the sled was set to 100 cm/s. If the steering wheel angle encoded a higher sled speed, it was capped at this maximum speed. The sled stopped when the steering wheel angle fell within −2 to 2 deg from the start angle, or when the steering wheel angle fell within −6 to 6 deg from the start angle and remained constant for 100 ms or started rising again (stopped steering prematurely or started a new steering movement). If the sled reached one of the ends of the track, it also stopped. Figure 1C shows the sled position, velocity and acceleration as a function of time for five representative trials. White noise was played via the earphones during the steering movement to mask auditory cues induced by the sled motion.

After the sled stopped, participants received feedback about their performance. First, both the current location of the body and the target location were presented on the screen for 1 s. This informed participants about how far they ended up from the target location and whether they undershot or overshot the target location. To encourage participants to be as accurate as possible, participants received “hit” feedback if the distance between the current location of the body and the target location was smaller than 4.5 cm. This “hit” area was represented on the screen by a translucent orange rectangular area (width 9 cm and height 25.4 cm) horizontally centred on the target location (Feedback I in Fig. 1B). After this, two horizontal feedback bars (width 15.2 cm and height 1 cm) were shown for 2 s (Feedback II in Fig. 1B). The centre of the feedback bars was green, flanked by orange and red areas towards the edges. A white bar on the upper feedback bar reiterated the displacement error, with the centre of the green area corresponding to the target location, and the left and right edges of the green area corresponding to an undershoot and overshoot of 4.5 cm, respectively (i.e., the “hit” window). A cheerful sound was played via the earphones if the participant “hit” the target. A white bar on the lower feedback bar showed the movement duration. Participants were encouraged to finish their steering movement within 800-1200 ms from movement start to ensure suprathreshold vestibular stimulation while remaining below the maximum sled speed. The centre of the green area of the feedback bar corresponded to a movement duration of 1000 ms, and the left and right edges of the green area corresponded to a movement duration of 800 and 1200 ms, respectively. If the displacement error or the movement duration was out of bounds (i.e., actual location of the white bar was more extreme than the left and right outer edges of the red areas of the feedback bar, corresponding to movement durations of 200 ms and 1800 ms, respectively), the white bar was presented on the outer edge of the feedback bar closest to the true location.

The next trial started after the feedback had disappeared. If the location of the sled at the end of the trial restricted its motion on the next trial to less than 45 cm, the sled was first passively moved to a new starting location. This starting location was 15 cm away from the middle of the sled track in the direction opposite of the upcoming target location, leaving approximately 60 cm for the upcoming displacement.

As described above, the steering gain (i.e., the gain between the angle of the steering wheel and the velocity of the sled) changed throughout the experiment. All participants experienced three different conditions: a random walk condition, a white noise condition, and a constant gain condition (Fig. 1D). In total, the experiment consisted of nine trial blocks, with three trial blocks per condition. Each trial block started with 36 trials specific to the condition. On the last of these condition-specific trials, the steering gain was always 1.0 cm/s per deg (baseline trial), and this trial was always followed by four trials with a high gain of 1.4 cm/s per deg (step trials) and six trials with the baseline gain of 1.0 cm/s per deg (washout trials).

For the random walk condition, the gains of the other condition-specific trials were generated backwards, starting from the baseline trial, in the following way:

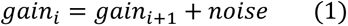

in which *i* is the trial number. Noise samples were drawn from a Gaussian distribution with a mean of 0 cm/s per deg and a standard deviation of 0.1 cm/s per deg. Random walks were drawn until a walk (excluding the baseline trial) met the following criteria: a mean gain between 0.99 and 1.01 cm/s per deg, a standard deviation between 0.139 and 0.141 cm/s per deg, and a lag-1 autocorrelation value higher than 0.8 (i.e., high predictability). Autocorrelation values were computed by dividing the autocovariance values by the variance of the gains, such that the autocorrelation values fell within −1 to 1. We controlled the standard deviation to ensure spread in the gains while avoiding gains more extreme than the gain on the step trials. The procedure was repeated three times per participant, yielding three random walks per participant.

For the white noise condition, the gains from the three random walks, except for the baseline trials, were shuffled. This was done 10,000 times per walk, and for each walk the instance with the lowest absolute lag-1 autocorrelation value was selected (all between 0.001 and −0.001). This way, the condition-specific trials in the white noise trial blocks had the same means and standard deviations as the condition-specific trials in the random walk trial blocks. In the constant gain condition, all 36 condition-specific trials in the three trial blocks had a gain of 1.0 cm/s per deg (baseline gain).

The conditions were presented in a random order per participant. The number of repetitions of all six possible combinations was balanced across participants whose data was included in the analysis (see below). The three trial blocks per condition were presented consecutively but in random order. At the end of each trial block, the percentage of “hit” trials was presented on the screen, followed by a short break (> 45 seconds) during which the lights in the experimental room were turned on to prevent dark adaptation. Before the experiment, all participants completed 18 practice trials with a baseline gain of 1.0 cm/s per deg, during which the experimenter was present for task instructions. In total, each participant completed 432 trials.

### Data analysis

Data were processed offline in MATLAB (v.R2017a; the MathWorks, Inc., Natick, MA). Data from two participants were excluded from the analysis because of their relatively low scores (average percentage of “hit” trials across trial blocks 48 and 58%; range included participants 66-90%). Trials during which participants rotated the steering wheel less than 7.5 deg or displaced the sled in the direction opposite of the target were excluded from the analysis. Additionally, trials during which the speed encoded by the steering wheel angle reached the set maximum of 100 cm/s or during which the sled reached one of the ends of the sled track were excluded. On average, one trial was excluded per participant (range 0-2 trials).

Movement onset was defined as the first time point that the steering wheel was rotated more than 2 deg. Movement end was defined as the first time point after movement onset that the steering wheel angle fell within −2 to 2 deg, or as the time point after which the steering wheel angle remained constant for at least 100 ms or reached a local minimum between −7.5 and 7.5 deg (i.e., failed to bring the steering wheel back to the start position or started a new steering movement). Movement duration was defined as the time between movement onset and movement end. Displacement error was defined as the difference between the location of the body at movement end and the target location. Negative errors represent undershoots and positive errors represent overshoots.

### Trial series regression analysis

To examine whether the predictability of the gains affected the steering behaviour on the condition-specific trials in the white noise and random walk condition, we performed a trial series regression analysis. For each time point *t* within trial *i*, we modelled the steering wheel angle *a* as a linear combination of a constant representing the average steering wheel angle at time point *t* across trials, the gain on trial *i* (i.e., the current trial), the gain on trial *i*-1 (i.e., the previous trial), and residual error *ε*:

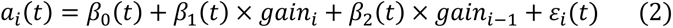

First, we selected the condition-specific trials (including the baseline trial but excluding the first trial of each trial block) and sampled the absolute steering wheel angle every 20 ms from 0 to 800 ms after movement onset for computational efficiency. Gains were z-scored based on the means and standard deviations of the gains on the included trials within the corresponding trial block. Trial blocks from the constant gain condition were not included in this analysis, because the gain was kept constant throughout the condition-specific trials, making it impossible to use a regression analysis. The regression model was fitted per sampled time point within a trial, per trial block of the white noise and random walk condition, and per participant, yielding 5,904 runs in total (41 time points x 6 trial blocks x 24 participants).

To check whether the autocorrelation in the gains in the random walk condition could potentially lead to autocorrelated regression coefficients, we used simulations of the regression model in Equation 2. These simulations showed that the regression coefficients could be reliably estimated, both when the autocorrelation values of the predictors were high, similar to the random walk condition, and when the autocorrelation values were close to zero, similar to the white noise condition. This shows that the gains were distinct enough across trials to reliably estimate the regression coefficients described in Equation 2 in both the white noise and the random walk condition.

Based on the regression coefficients from the regression fits, we additionally predicted the absolute steering wheel angle as a function of time, as well as the separate effects of the current and the previous gain on the steering wheel angle, for the baseline and step trials. To compute the predictions, gains were z-scored based on the means and standard deviations of the gains from the included condition-specific trials within the corresponding trial block and were multiplied with the regression coefficients following Equation 2. Predictions were made per sampled time point within a trial, per trial block of the white noise and random walk condition, and per participant. If the predicted steering wheel angle fell below 2 deg, the steering movement ended.

### Statistics

Statistical analyses were done in MATLAB and R (v.4.0.1; see R Core Team, 2017). The alpha value for statistical significance was set to .05, and this value was Bonferroni-corrected in case of multiple comparisons (exact value of alpha specified with the results of the tests). To compare the overall performance across conditions and trial block repetitions, we examined the average displacement error, movement duration and maximum absolute steering wheel angle across trials within a block using a two-way repeated-measures ANOVA with condition (white noise, random walk, and constant gain) and trial block number (first, second, and third repetition) as within-subject factors using the ez-package in R (v.4.4-0; see Lawrence, 2016). The results were adjusted according to the Greenhouse-Geisser correction in case of violations of sphericity, and we report the generalized eta-squared (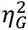) as a measure of the effect size. To examine the responses to the step changes in the gain across conditions, we averaged the displacement error, movement duration and maximum absolute steering wheel angle on the baseline trial and the step trials across trial blocks within a condition and examined differences between the trials and conditions using a two-way repeated-measures ANOVA with condition (white noise, random walk, and constant gain) and trial (baseline and first, second, third, and fourth step trial) as within-subject factors. We used paired-samples t-tests to directly compare the groups post hoc. To compare the results of the regression fits across the white noise and random walk condition, we averaged the regression coefficients across trial blocks within a condition and compared the values of the regression coefficients across the two conditions at each time point with a paired-samples t-test in MATLAB.

## Results

We used a closed-loop steering experiment in which participants steered a linear sled to align their body with a memorized visual target. We varied the steering gain in three different ways, and examined whether participants took the predictability of the gain into account in their steering behaviour (internal model hypothesis) or whether participants simply integrated the vestibular information over time (path integration hypothesis).

### General observations

Figure 2A shows the average displacement error across participants as a function of trial number for each of the three conditions. Participants hit the target on average in 75% of trials (range 66-90%). The average displacement error was close to zero in all three conditions (white noise: *M* = 0.25 cm, *SD* = 1.12 cm; random walk: *M* = 0.03 cm, *SD* = 1.06 cm; constant gain: *M* = 0.30 cm, *SD* = 1.12 cm) and in all three trial blocks within a condition (first repetition: *M* = 0.18 cm, *SD* = 1.22 cm; second repetition: *M* = 0.32 cm, *SD* = 1.05 cm; third repetition: *M* = 0.09 cm, *SD* = 1.02 cm). In line, the overall displacement error did not differ significantly across the conditions (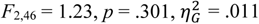) or across the three trial blocks within a condition (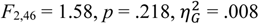). Additionally, there was no significant interaction effect between the condition and the trial block number (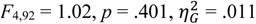). The included participants were thus able to hit the target in most trials.

**Figure 2.**
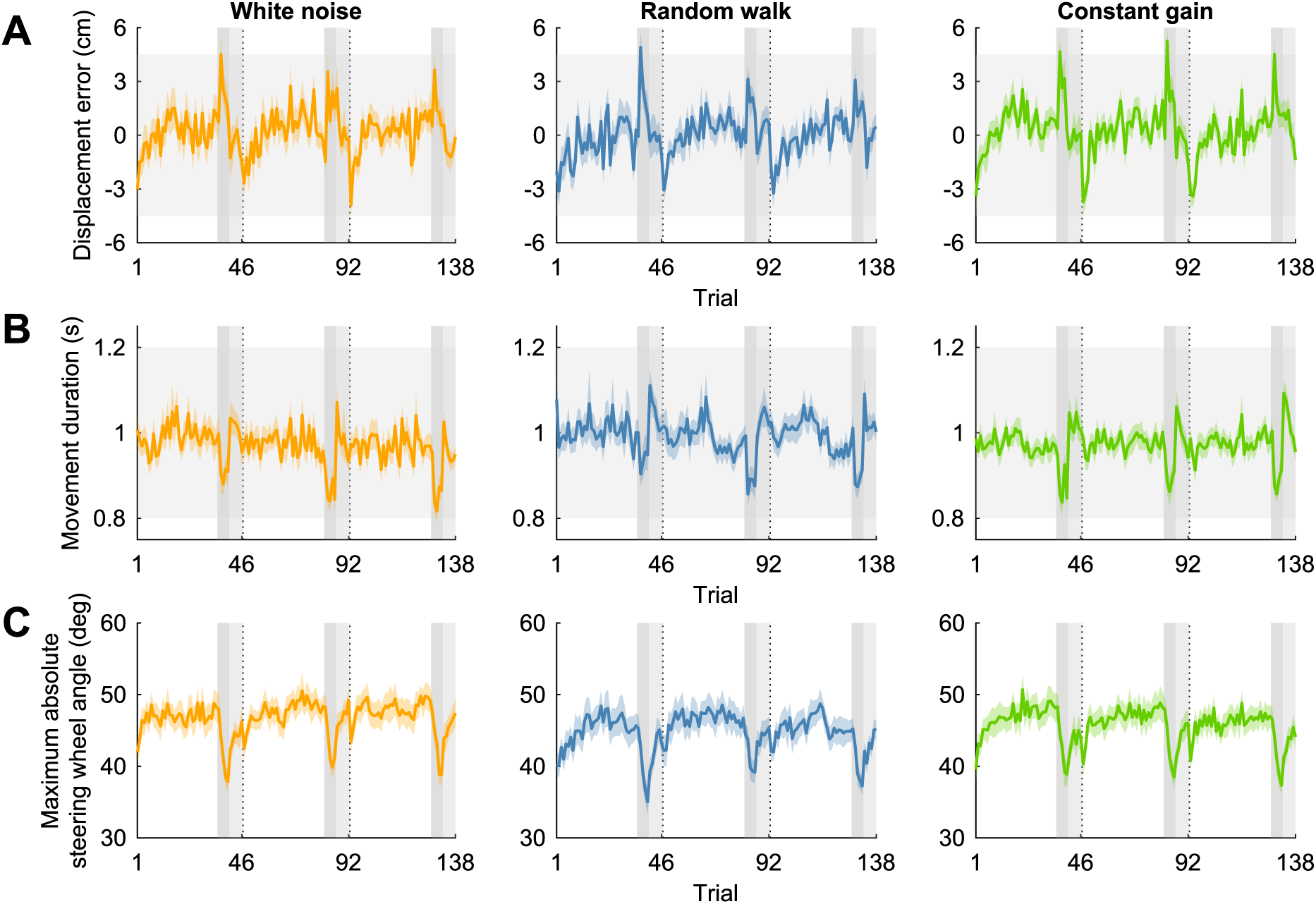
Displacement error, movement duration and maximum absolute steering wheel angle. A) Mean displacement error across participants as a function of trial number grouped based on the experimental condition (panels). Negative numbers represent undershoots; positive numbers represent overshoots. Coloured shaded areas represent between-subjects means ± SE. Participants completed three trial blocks per condition in sequence and the conditions were presented in a random order per participant. Each trial block was concluded with a baseline trial, followed by four step trials with a high gain (dark grey vertical areas) and six washout trials with the baseline gain (light grey vertical areas). Dashed vertical lines represent breaks and horizontal light grey bands show the range of displacement errors within which participants “hit” the target. B) Same configuration as in A, but with the mean movement duration across participants. Horizontal light grey bands show the time window within which participants were encouraged to finish their steering movement. C) Same configuration as in A, but with the mean maximum absolute steering wheel angle across participants.

Figure 2B shows the average movement duration in the same format as in Figure 2A. On average, participants finished their steering movement within the imposed time window from 800 to 1200 ms in all three conditions (white noise: *M* = 970 ms, *SD* = 97 ms; random walk: *M* = 994 ms, *SD* = 108 ms; constant gain: *M* = 972 ms, *SD* = 86 ms) and in all three trial blocks (first repetition: *M* = 987 ms, *SD* = 101 ms; second repetition: *M* = 974 ms, *SD* = 99 ms; third repetition: *M* = 975 ms, *SD* = 93 ms). There was no significant difference in the overall movement duration across conditions (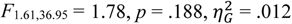) or trial blocks (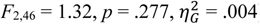). Additionally, there was no significant interaction effect between the condition and the trial block number (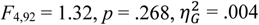).

Figure 2C shows the average maximum absolute steering wheel angle in the same format as in Figure 2A. The average maximum absolute steering wheel angle was similar across conditions (white noise: *M* = 46.7 deg, *SD* = 7.0 deg; random walk: *M* = 44.9 deg, *SD* = 6.9 deg; constant gain: *M* = 45.7 deg, *SD* = 7.0 deg) and repetitions (first repetition: *M* = 45.4 deg, *SD* = 7.1 deg; second repetition: *M* = 46.2 deg, *SD* = 7.0 deg; third repetition: *M* = 45.7 deg, *SD* = 6.9 deg). In line, the maximum absolute steering wheel angle was not significantly affected by the condition (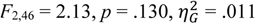) or the trial block number (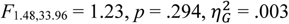), nor was there an interaction effect (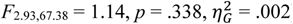).

### Trial series regression analysis

These findings suggest that, overall, the steering behaviour was similar across conditions and trial blocks. However, in the white noise and random walk condition, the steering gain varied from trial to trial and across participants. Averaging across trials within a trial block and across participants could therefore mask effects of the gain on the steering behaviour at the single-trial level. To examine the relationship between the gain and the steering behaviour at the single-trial level we fitted a trial series regression model (see Methods). Using this approach, we describe for each condition-specific trial of the white noise and random walk condition the steering wheel angle at a certain point in time as a function of the gain on the current trial, the gain on the previous trial, and a constant (or offset). Figure 3 shows the results of the regression model. The regression coefficient representing the constant as a function of time follows the average steering wheel profile (Fig. 3A). The constant did not differ significantly across the two conditions (smallest p-value: *p* = .113 at 560 ms after movement onset; Bonferroni-corrected *α* = .0012).

**Figure 3.**
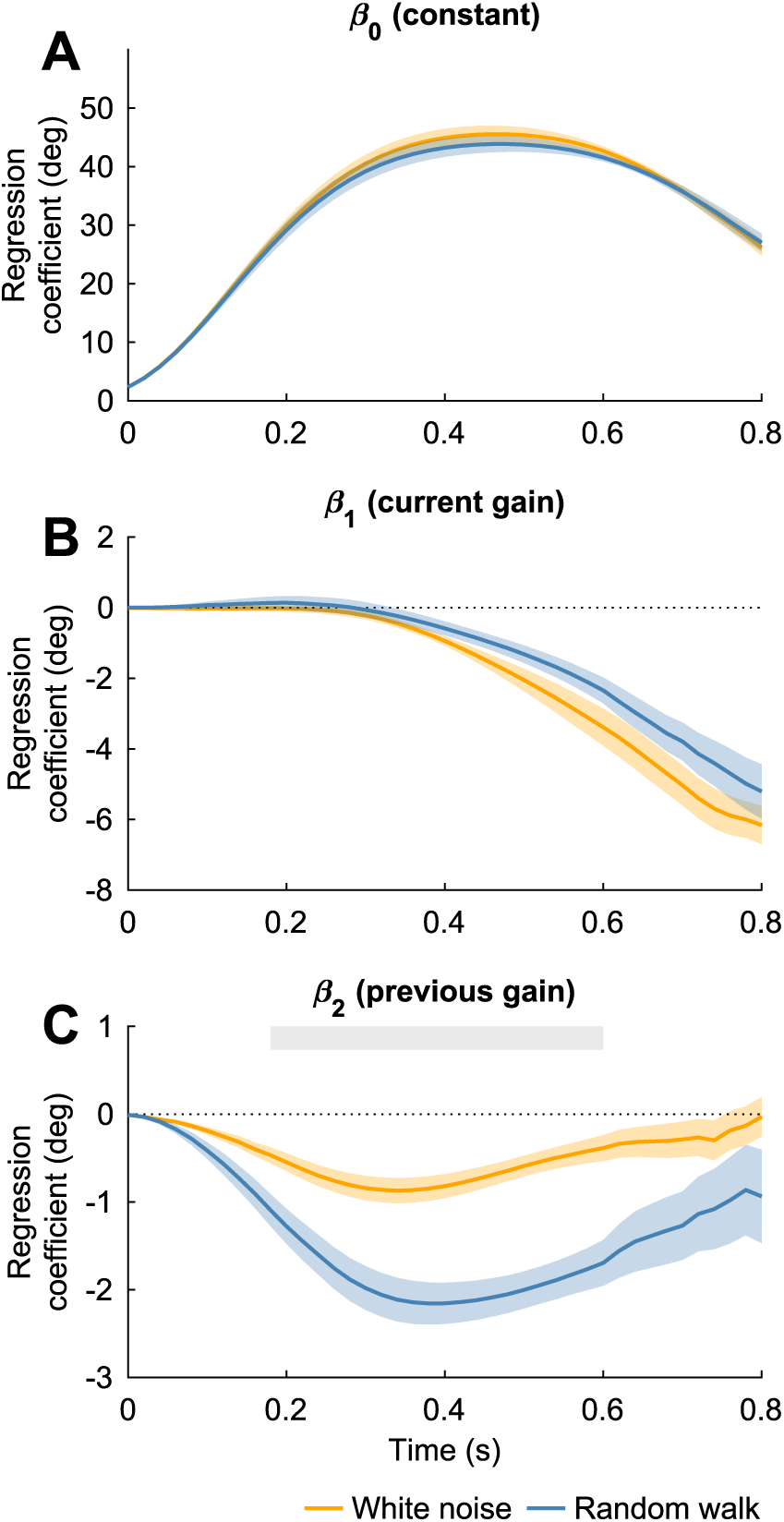
Trial series regression model. The trial series regression model described the steering wheel angle as a function of a constant, the gain on the current trial, and the gain on the previous trial. A) Mean value of the regression coefficient representing the constant across trial blocks and participants as a function of time grouped based on experimental condition (coloured lines). Values represent the average steering wheel angle as a function of time across the condition-specific trials within a trial block. Coloured shaded areas represent between-subjects means ± SE. B) Same configuration as in A, but with the mean value of the regression coefficient of the current gain across trial blocks and participants. Negative regression coefficients indicate that participants decreased and increased the steering wheel angle with an increase and decrease in the gain relative to the mean, respectively. C) Same configuration as in A, but with the mean value of the regression coefficient of the previous gain across trial blocks and participants. The regression coefficient differed significantly between the conditions from 180 to 600 ms after movement onset, as also indicated by the light grey shaded horizontal area (*p* < .0012).

Figure 3B shows the regression coefficient for the current gain, which was zero at the beginning of the steering movement and started to decrease after approximately 300 ms in the white noise condition (significantly different from zero from 340 to 800 ms after movement onset; range p-values from *p* = .0008 to *p* < .0001; Bonferroni-corrected *α* = .0012) as well as in the random walk condition (significantly different from zero from 440 to 800 ms after movement onset; range p-values from *p* = .0004 to *p* < .0001; Bonferroni-corrected *α* = .0012). Negative regression coefficients indicate that participants decreased the steering wheel angle with an increase in the gain relative to the mean gain and vice versa. Participants thus reacted adequately to changes in the gain from trial to trial by steering against the gain change midway the steering movement. The regression coefficient for the current gain did not differ across the two conditions (smallest p-value: *p* = .0074 at 540 ms after movement onset; Bonferroni-corrected *α* = .0012).

Figure 3C shows the regression coefficient for the previous gain, which decreased almost immediately after movement onset and started increasing again after approximately 400 ms in both conditions. The regression coefficient differed significantly from zero from 40 to 540 ms in the white noise condition (range p-values from *p* = .0011 to *p* < .0001; Bonferroni-corrected *α* = .0012), and from 80 to 660 ms in the random walk condition (range p-values from *p* = .0009 to *p* < .0001; Bonferroni-corrected *α* = .0012). The effect was small in the white noise condition, as expected. The regression coefficient was significantly more negative in the random walk condition than in the white noise condition (significant from 180 to 600 ms after movement onset; range p-values from *p* = .0012 to *p* < .0001; Bonferroni-corrected *α* = .0012), indicating that the effect of the previous gain on the steering wheel angle was larger in the random walk condition. This is in line with the internal model hypothesis, as it is more advantageous to take the previous gain into account in the random walk condition because it is more predictive of the current gain due to the high autocorrelation.

### Step trial analysis

To examine whether these differences in the steering strategy across the conditions were also directly visible in the steering behaviour after larger jumps in the gain, we added four step trials with a high gain of 1.4 cm/s per deg to the end of each trial block. All step trials were preceded by a baseline trial and were followed by six washout trials, all with a gain of 1.0 cm/s per deg.

Figure 4 shows the displacement error, movement duration and maximum absolute steering wheel angle for the baseline and step trials, grouped based on the condition and averaged across trial blocks and participants. To examine participants’ responses to the step changes in the gain, we compared the baseline trial and the step trials across conditions. In all conditions, the average displacement error on the first step trial was positive and larger than the average displacement error on the baseline trial (Fig. 4A). There was a significant main effect of the trial on the displacement error (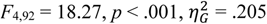), and post hoc paired-samples t-tests revealed that there was a significant difference between the baseline trial and all four step trials (range p-values from *p* = .002 to *p* < .0001; Bonferroni-corrected *α* = .005), and between the first step trial and the subsequent step trials (all p-values < .0001; Bonferroni-corrected *α* = .005). There was no significant main effect of the condition on the displacement error (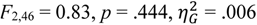), or a significant interaction effect (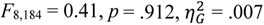). However, in all three conditions the overshoot of the target location on the first step trial was smaller than 12 cm, which is the displacement error that would be expected if participants did not respond to the increase in the gain (target distance of 30 cm and gain increase from 1.0 cm/s per deg to 1.4 cm/s per deg). This suggests that participants changed their steering movement online during the first step trial to compensate for the increase in the gain in all three conditions.

**Figure 4.**
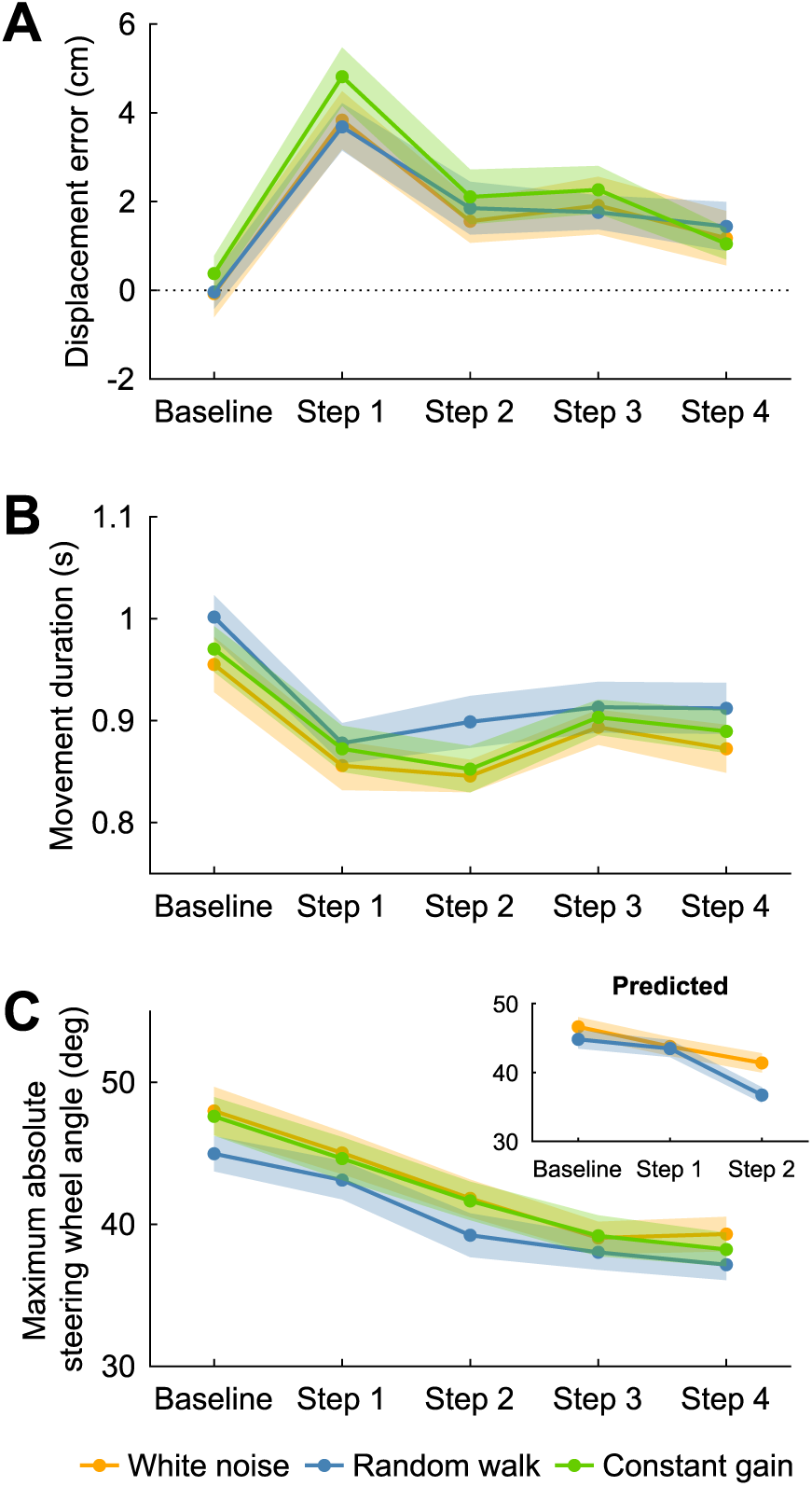
Displacement error, movement duration and maximum absolute steering wheel angle on baseline and step trials. A) Mean displacement error across trial blocks and participants as a function of the trial grouped based on the experimental condition (coloured lines). Negative numbers represent undershoots; positive numbers represent overshoots. Coloured shaded areas represent between-subjects means ± SE. B) Same configuration as in A, but with the mean movement duration across trial blocks and participants. C) Same configuration as in A, but with the mean maximum absolute steering wheel angle across trial blocks and participants. Inset shows the maximum absolute steering wheel angles for the baseline and first and second step trials of the white noise and random walk conditions, predicted based on the results of the trial series regression model.

This is confirmed by changes in the movement duration (Fig. 4B) and maximum absolute steering wheel angle (Fig. 4C) from the baseline trial to the step trials. The movement duration differed significantly across trials in all conditions (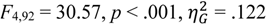), with significantly shorter movement durations on the step trials than on the baseline trial (all p-values < .0001; Bonferroni-corrected *α* = .005). Interestingly, the movement duration increased again across the step trials, with significantly longer movement durations on the third step trial than on the first and second step trials (range p-values from *p* = .0001 to *p* < .0001; Bonferroni-corrected *α* = .005). There was no significant main effect of the condition on the movement duration (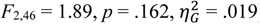), or an interaction effect (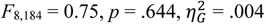). Similarly, the maximum absolute steering wheel angle differed significantly across trials in all conditions (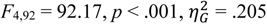), with a significantly larger maximum absolute steering wheel angle on the baseline trial than on all four step trials (all p-values < .001; Bonferroni-corrected *α* = .005). The maximum absolute steering wheel angle continued to decrease significantly across the first three step trials (all p-values < .001; Bonferroni-corrected *α* = .005). There was no significant main effect of the condition on the maximum absolute steering wheel angle (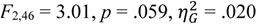), or an interaction effect between the trial and the condition (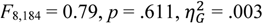). Participants seemed to fine-tune their steering behaviour after the large jump in the gain by increasing the movement duration again slightly and continuing to decrease the maximum absolute steering wheel angle across the step trials, thereby minimizing the displacement error while simultaneously adhering to the imposed time window.

The similarity in the correction across conditions and the fine-tuning of the steering behaviour across the step trials is also shown in Figure 5. In this figure, the steering wheel angle as a function of time is normalized relative to the baseline trials. For each participant and trial block, we first resampled the steering wheel angles of the baseline and the four step trials to 200 samples per trial using linear interpolation. The movement duration and steering wheel angles on these trials were then normalized by dividing them by the movement duration and the maximum absolute steering wheel angle of the corresponding baseline trial, respectively. Normalized steering wheel angles were averaged across trial blocks and participants, and a corresponding linearly spaced time vector of 200 samples was created for each trial running from zero, representing movement onset, to the mean normalized movement duration across trial blocks and participants.

**Figure 5.**
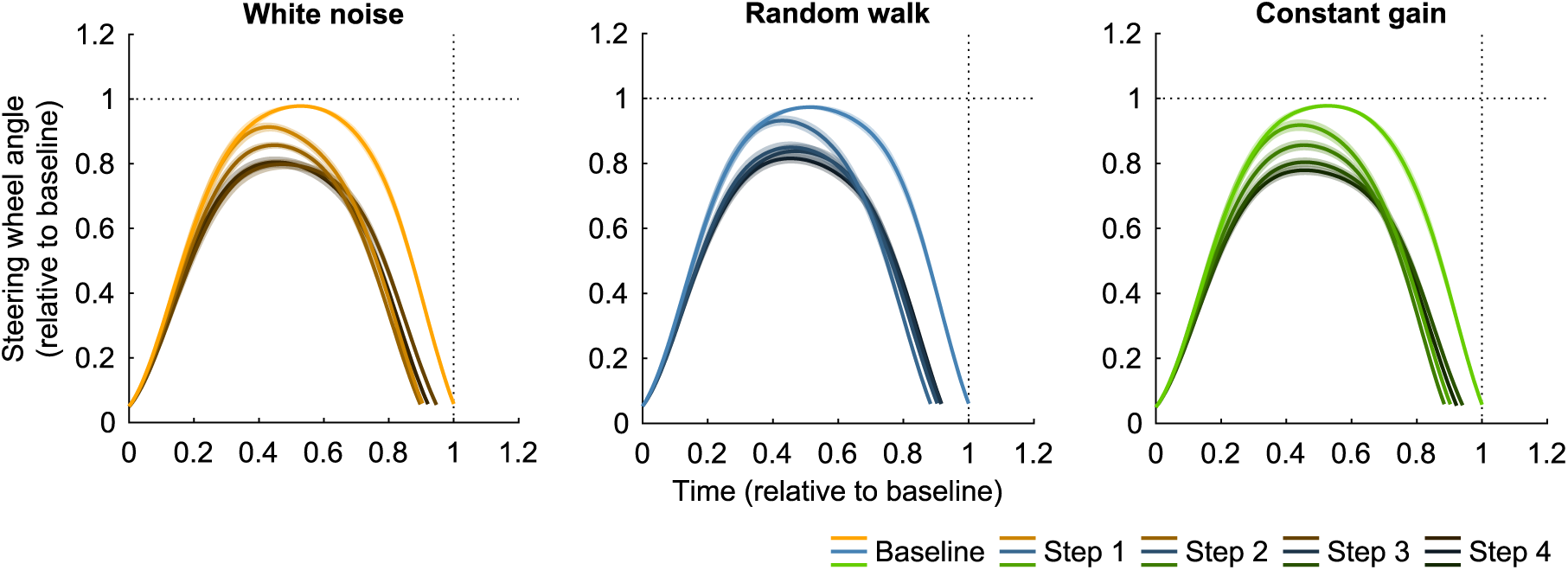
Steering behaviour on the baseline and step trials. Average absolute steering wheel angle as a function of time across trial blocks and participants for the baseline and step trials grouped based on the experimental condition (panels). Values were normalized relative to baseline. Coloured shaded areas represent between-subjects means ± SE.

As described above, participants decreased both the movement duration and the maximum absolute steering wheel angle in response to the increase in the gain from the baseline to the first step trial. They did this already early on within the first step trial. Even though the responses to the higher gain are similar across conditions, the decrease in the maximum absolute steering wheel angle from the baseline trial to the first step trial seems to be slightly smaller in the random walk condition, as also shown in Figure 4C. Participants continued to decrease the maximum absolute steering wheel angle across the subsequent step trials, while slightly increasing the movement duration again towards the baseline movement duration.

Based on the results of the regression model fitted to the condition-specific trials, we made predictions for the steering wheel angle as a function of time for the baseline and the first two step trials in the white noise and random walk condition. Figure 6A shows the mean experimentally observed absolute steering wheel angles as a function of time for the baseline and the first two step trials. These steering wheel profiles are the same as the profiles shown in Figure 5, but without the baseline normalization. Overall, the steering wheel angles were slightly smaller in the random walk condition than in the white noise condition, and this difference is accurately predicted based on the regression model, as shown in the right panel in Figure 6A.

**Figure 6.**
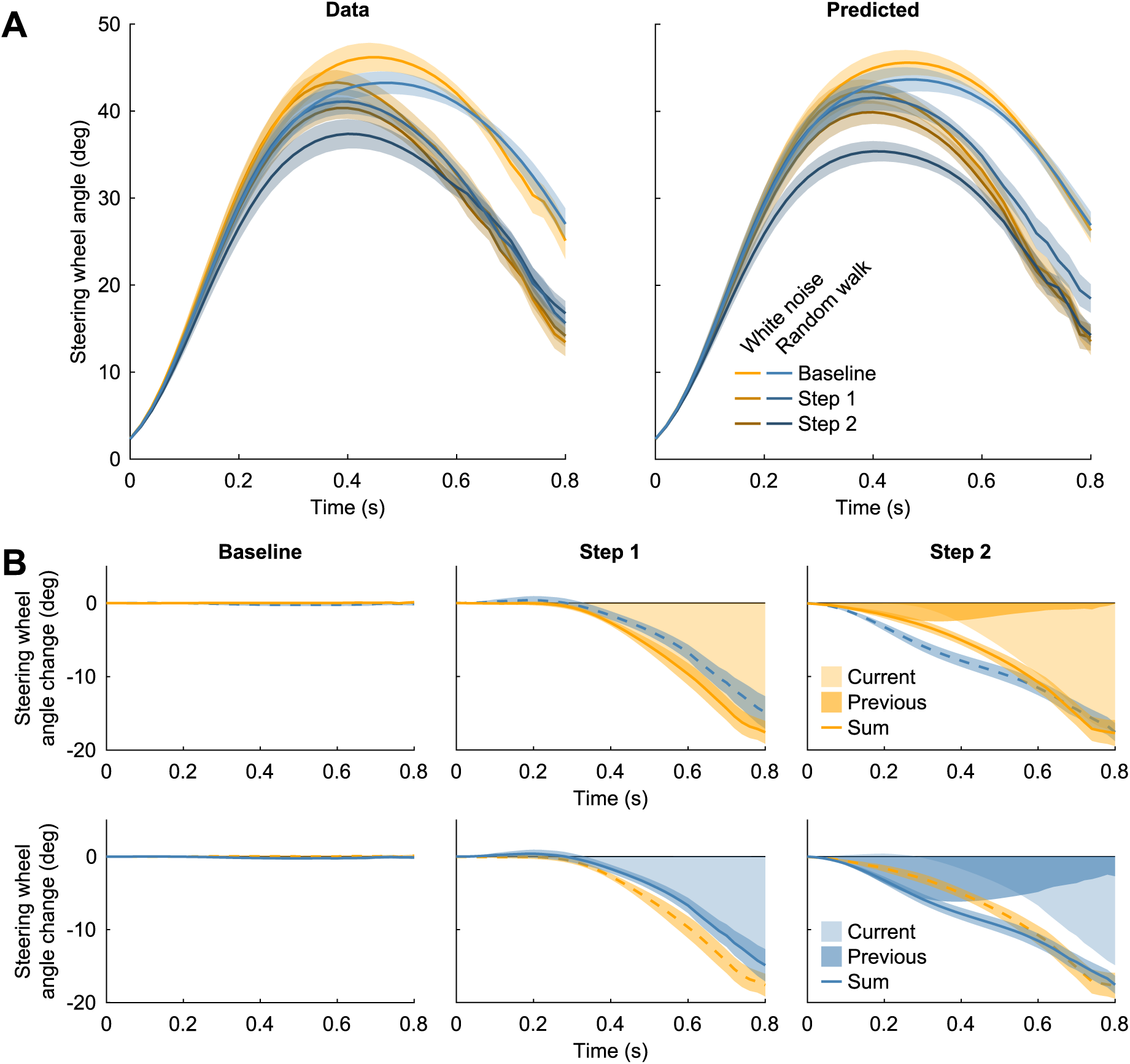
Predicted steering behaviour on the baseline and step trials, based on the trial series regression model. A) Mean absolute steering wheel angle as a function of time across trial blocks and participants for the baseline and the first two step trials in the white noise and random walk conditions (left panel), and the predicted values based on the regression coefficients of the regression model (right panel). Steering wheel angles were predicted for the baseline trial and the first two step trials, per trial block and participant. Coloured shaded areas represent between-subjects means ± SE. B) Mean predicted change in the steering wheel angle relative to the average steering wheel profile, represented by the constant of the regression model, based on the current and previous gain (shaded areas) as well as the sum (solid coloured lines, coloured shaded areas represent between-subjects means ± SE) for the baseline and the first two step trials (horizontal panels) of the white noise and random walk conditions (upper and lower panels, respectively) across trial blocks and participants. Dashed coloured lines show the sum of the predicted change in the same trial for the other condition as a reference line.

We could additionally separate the effects of the current and the previous gain on the changes in the steering wheel profiles across the baseline and the first two step trials. Figure 6B shows the products of the regression coefficients for the current and the previous gain and the z-scored gains as a function of time for each of the three trials. For the baseline trials, the z-scored current gain was close to zero, as the gain on the baseline trial was always 1.0 cm/s per deg and the mean gain across the condition-specific trials within a trial block was set to be close to the baseline gain. Additionally, since the gain on the trial before the baseline trial varied across trial blocks and participants, the average contribution of the previous gain is also close to zero for the baseline trial. The steering wheel angle as a function of time on this trial was thus similar to the constant in the regression model, representing the average steering wheel angle as a function of time across the condition-specific trials within the trial block.

On the first step trial, the effect of the gain experienced on the previous trial, the baseline trial, on the steering wheel angle is again very close to zero. The effect of the gain on the current trial is however large in both conditions. The predicted change in the steering wheel angle relative to the average angle is slightly greater in the white noise condition due to the slightly more negative value for regression coefficient *β_1_*, as shown in Figure 3B. On the second step trial, there is an effect of the gain on both the previous and the current trial on the steering wheel angle. In the random walk condition, the effect of the previous gain is larger than in the white noise condition due to a significantly more negative value for regression coefficient *β_2_*, as shown in Figure 3C. This leads to a greater overall reduction in the steering wheel angle over time relative to the mean angle, starting rather early on in the movement. This can also be observed, albeit a little less pronounced, from the greater reduction in the maximum absolute steering wheel angle in the random walk condition from the first to the second step trial in Figure 4C (see the inset for the predicted maximum absolute steering wheel angle), and the greater difference between the steering wheel profiles of the first and the second step trial in the random walk condition in Figure 5. Hence, there are differences in the steering behaviour across the white noise and random walk conditions, which can be mainly observed on the second step trial, due to different effects of the gain on the previous and current trial on the steering wheel angle.

## Discussion

In this study, participants used a steering wheel to move their body to a memorized visual target location. They were exposed to three experimental conditions, in which the gain between the steering wheel angle and the velocity of the linear motion platform varied with different levels of predictability from one trial to the next. In the white noise condition, the steering gain varied randomly from trial to trial (i.e., not predictable), in the random walk condition it was moderately predictable, and in the constant gain condition it remained constant across trials (i.e., highly predictable). The goal was to examine whether participants took the predictability of the gain into account in their steering behaviour, by forming and relying on an internal model of the steering dynamics, or whether they simply relied on the vestibular feedback in their steering, as in path integration.

Using a trial series regression analysis, we have shown that participants used a different steering strategy for the white noise and random walk conditions (Fig. 3). The average steering wheel angle and the regression coefficient for the current gain were similar across conditions throughout the trial, but the regression coefficient for the previous gain was significantly more negative in the random walk condition from 180 to 600 ms after movement onset. This suggests that participants decreased the steering wheel angle more in the random walk condition than in the white noise condition if the gain on the previous trial was higher than the average gain, and vice versa. Based on the results of the regression model, we also predicted the subtle differences between the white noise and random walk condition in the changes in the steering behaviour from the baseline to step trials, in which the gain was suddenly higher for four consecutive trials (Fig. 6). Participants thus took the previous gain into account in the random walk condition, which is a useful strategy given the high autocorrelation in the gains. We conclude that participants formed an internal representation of the steering dynamics, which is in line with the internal model hypothesis.

In all conditions, including the constant gain condition, participants decreased the maximum absolute steering wheel angle and the movement duration on the first step trial. Across the subsequent step trials (Fig. 5), participants improved their adaptation to the new steering dynamics by simultaneously increasing the movement duration and decreasing the maximum absolute steering wheel angle to adhere to the time window imposed in the experiment, similar as in van Helvert et al. (2022). These tactful changes in the steering behaviour underline the idea that participants built and updated an internal model of the steering dynamics and the associated self-motion based on the vestibular feedback. In principle, we could have also predicted the steering behaviour on the third and fourth step trial based on the fitted regression model. For these trials, both the gain on the current and the previous trial would be the same as for the second step trial, and the prediction would thus be that the steering behaviour remains the same across these three trials. Even though the changes in the steering behaviour are relatively small across these step trials, see for example Figure 4 and 5, it seems plausible that participants revised their steering strategy on these trials, given that the dynamics on the step trials were different from the dynamics experienced during the condition-specific trials.

In our previous study (van Helvert *et al*., 2022), participants performed a similar steering experiment but the steering gain changed only twice during the whole experiment, comparable with the constant gain condition in the current experiment. We found that participants responded rapidly to these changes in the steering dynamics, suggesting that participants had some expectations about their velocity and the steering dynamics, but we could not further distinguish between the internal model hypothesis and the path integration hypothesis. Here, we dissociate the contribution of vestibular feedback and predictions by changing the steering dynamics across trials with different levels of predictability. We show that participants use the vestibular feedback during the trial to estimate their self-motion, but also that their steering behaviour depends on the predictability of the steering gain. Important to note, this conclusion is based mainly on the results of the regression model, in which the data from the constant gain condition could not be included. Due to the high predictability of the steering gain, we expected participants to respond slowest on the step trials in the constant gain condition, but the data did not support this notion. To study steering behaviour with a high predictability of the steering gain, future studies could include white noise and random walk conditions with varying levels of variability, similar to Burge et al. (2008) who studied the trade-off between prediction and estimation based on sensory feedback in reach adaptation.

Stavropoulos et al. (2022) also used a closed-loop steering experiment to study the role of vestibular feedback and predictions in self-motion estimation. Participants used a joystick to navigate to a target while the steering dynamics changed from trial to trial following a bounded random walk. They found that the steering behaviour was biased with responsive steering control and concluded that participants were not able to accurately steer and build an internal model of the steering dynamics based on the vestibular feedback alone. Our previous results suggested that participants can accurately estimate their self-motion and suggest that they build an internal model of the steering dynamics based on just vestibular feedback (van Helvert *et al*., 2022), and here we show that they can even do this under steering dynamics that change from trial to trial. This discrepancy between the results might be explained by the fact that the velocity of the motion platform used by Stavropoulos et al. (2022) was close to constant. This may have made it more difficult for participants to estimate their self-motion, as the vestibular organs and more specifically the otoliths, which process information about translational motion, are known to be mainly sensitive to acceleration (Fernandez & Goldberg, 1976; Benson *et al*., 1986; Fitze *et al*., 2023). Also, participants in our experiments received feedback about their performance at the end of each trial, which is likely to have removed any possible biases in participants’ self-motion estimates.

The present study is based on previous studies of visuomotor and force field adaptation in reaching movements that examined the role of sensory feedback and predictions (Burge *et al*., 2008; Wei & Körding, 2010; Gonzalez Castro *et al*., 2014). These studies showed that participants respond faster to perturbations if the mapping between the reaching movement and the sensory feedback is more uncertain, due to a greater reliance on the feedback. Additionally, Gonzalez Castro et al. (2014) examined motor adaptation when the force field perturbation strength varied randomly across trials and when it varied according to a random walk. They showed that participants relied more on predictions of the perturbation in the random walk condition. This is in line with our finding that the effect of the previous gain on the steering behaviour is more pronounced in the random walk condition than in the white noise condition.

Our results suggest that participants build an internal model of the steering dynamics to estimate their self-motion during active steering. Multiple studies have looked for neural markers of such an internal model and have tried to unravel its location in the vestibular processing pathway (Roy & Cullen, 2001; Page & Duffy, 2008; Egger & Britten, 2013; Jacob & Duffy, 2015; Lakshminarasimhan *et al*., 2023). In general, the cerebellum is thought to play an important role in the internal model computations for self-motion estimation (Brooks *et al*., 2015; Laurens & Angelaki, 2017; Rineau *et al*., 2023; Cullen, 2023), also because of its projections to the vestibular nuclei. Neurons in the vestibular nuclei are known to distinguish between active and passive self-motion, being less sensitive to actively generated, and thus predictable, self-motion (Cullen *et al*., 2011). However, these neurons respond similarly to passive self-motion and self-motion generated by a steering movement (Roy & Cullen, 2001). One explanation for this may be that the monkeys in the experiment were not trained enough to build an internal model of the steering dynamics. Another explanation may be that the cerebellum does not predict the sensory consequences of the steering movement, and that the internal model of the steering movement is located more downstream in the vestibular processing pathway (Alefantis *et al*., 2022). Similarly, during the processing of the visual reafference of steering movements in monkeys, markers for an internal model were found in the medial superior temporal area (Page & Duffy, 2008) and the posterior parietal cortex (Lakshminarasimhan *et al*., 2023).

The sensorimotor processes that underlie driving have gained additional interest with the development of automated vehicles (Nash *et al*., 2016; Nash & Cole, 2020). Nash and Cole (2016) have described these sensorimotor processes in detail, and have shown that a driver model that includes an internal model of the mapping between the steering wheel angle and the sensory feedback accurately describes human steering behaviour in their experimental set up (Nash & Cole, 2020). Our results are in line with these findings. Based on the predictions of such an internal model, feedforward control actions can be made, which can be extremely important in driving given the delays in the sensorimotor system (Nash *et al*., 2016). Along these lines, the present results may also stimulate novel concepts for artificial navigation systems, e.g., those providing independent mobility to sensory-deprived people and vehicle control.

In conclusion, our results show that participants take the predictability of changes in the steering dynamics into account during driving. This suggests that participants build an internal model of the gain between the steering wheel angle and their self-motion, and use this model to predict the vestibular reafference in driving and self-motion estimation.

## Additional information section

### Data availability statement

For review purposes, all data and code are available from the Radboud Data Repository via the following URL: di.dcc.DSC_2023.00037_462. Upon publication, all data and code will be made publicly available via the persistent identifier currently reserved for this collection: https://doi.org/10.34973/szbh-kc35.

### Author contributions

M.J.L.H., L.P.J.S., R.J.B. and W.P.M. conceived and designed research; M.J.L.H. performed experiments; M.J.L.H. analyzed data; M.J.L.H., L.P.J.S., R.J.B. and W.P.M. interpreted results of experiments; M.J.L.H. prepared figures; M.J.L.H., L.P.J.S., R.J.B. and W.P.M. drafted manuscript; M.J.L.H., L.P.J.S., R.J.B. and W.P.M. edited and revised manuscript. All authors approved the final version of the manuscript and agree to be accountable for all aspects of the work in ensuring that questions related to the accuracy or integrity of any part of the work are appropriately investigated and resolved.

## Acknowledgements

We would like to thank Brandon Rasman for his helpful comments on an earlier version of this manuscript.

## References

1. Alefantis P, Lakshminarasimhan KJ, Avila E, Noel J-P, Pitkow X & Angelaki DE (2022). Sensory evidence accumulation using optic flow in a naturalistic navigation task. J Neurosci 42, 5451– 5462.

2. Benson AJ, Spencer MB & Stott J (1986). Thresholds for the detection of the direction of whole-body, linear movement in the horizontal plane. Aviat Sp Environ Med 57, 1088–1096.

3. Britten KH (2008). Mechanisms of self-motion perception. Annu Rev Neurosci 31, 389–410.

4. Britton Z & Arshad Q (2019). Vestibular and multi-sensory influences upon self-motion perception and the consequences for human behavior. Front Neurol 10, 63.

5. Brooks JX, Carriot J & Cullen KE (2015). Learning to expect the unexpected: Rapid updating in primate cerebellum during voluntary self-motion. Nat Neurosci 18, 1310–1317.

6. Brooks JX & Cullen KE (2019). Predictive sensing: The role of motor signals in sensory processing. Biol Psychiatry Cogn Neurosci Neuroimaging 4, 842–850.

7. Burge J, Ernst MO & Banks MS (2008). The statistical determinants of adaptation rate in human reaching. J Vis 8, 20.

8. Campos JL, Butler JS & Bülthoff HH (2012). Multisensory integration in the estimation of walked distances. Exp Brain Res 218, 551–565.

9. Carriot J, Brooks JX & Cullen KE (2013). Multimodal integration of self-motion cues in the vestibular system: Active versus passive translations. J Neurosci 33, 19555–19566.

10. Cheng Z & Gu Y (2018). Vestibular system and self-motion. Front Cell Neurosci 12, 456.

11. Cullen KE (2023). Internal models of self-motion: Neural computations by the vestibular cerebellum. Trends Neurosci; DOI: 10.1016/J.TINS.2023.08.009.

12. Cullen KE, Brooks JX, Jamali M, Carriot J & Massot C (2011). Internal models of self-motion: Computations that suppress vestibular reafference in early vestibular processing. Exp Brain Res 210, 377–388.

13. DeAngelis G & Angelaki D (2011). Visual–vestibular integration for self-motion perception. In The Neural Bases of Multisensory Processes, ed. Murray MM & Wallace MT, pp. 629–650. CRC Press/Taylor & Francis. Available at: https://www.ncbi.nlm.nih.gov/books/NBK92839/ [Accessed September 18, 2023].

14. Egger SW & Britten KH (2013). Linking sensory neurons to visually guided behavior: Relating MST activity to steering in a virtual environment. Vis Neurosci 30, 315–330.

15. Fernandez C & Goldberg JM (1976). Physiology of peripheral neurons innervating otolith organs of the squirrel monkey. III. Response dynamics. J Neurophysiol 39, 996–1008.

16. Fitze DC, Mast FW & Ertl M (2023). Human vestibular perceptual thresholds — A systematic review of passive motion perception. Gait Posture 107, 83–95.

17. Genzel D, Firzlaff U, Wiegrebe L & MacNeilage PR (2016). Dependence of auditory spatial updating on vestibular, proprioceptive, and efference copy signals. J Neurophysiol 116, 765–775.

18. Glasauer S, Amorim MA, Viaud-Delmon I & Berthoz A (2002). Differential effects of labyrinthine dysfunction on distance and direction during blindfolded walking of a triangular path. Exp Brain Res 145, 489–497.

19. Gonzalez Castro LN, Hadjiosif AM, Hemphill MA & Smith MA (2014). Environmental consistency determines the rate of motor adaptation. Curr Biol 24, 1050–1061.

20. van Helvert MJL, Selen LPJ, van Beers RJ & Medendorp WP (2022). Predictive steering: Integration of artificial motor signals in self-motion estimation. J Neurophysiol 128, 1395–1408.

21. ter Horst AC, Koppen M, Selen LPJ & Medendorp WP (2015). Reliability-based weighting of visual and vestibular cues in displacement estimation ed. Ben Hamed S. PLoS One 10, e0145015.

22. Jacob MS & Duffy CJ (2015). Steering transforms the cortical representation of self-movement from direction to destination. J Neurosci 35, 16055–16063.

23. Kaski D, Quadir S, Nigmatullina Y, Malhotra PA, Bronstein AM & Seemungal BM (2016). Temporoparietal encoding of space and time during vestibular-guided orientation. Brain 139, 392–403.

24. Keshavarzi S, Velez-Fort M & Margrie TW (2023). Cortical integration of vestibular and visual cues for navigation, visual processing, and perception. Annu Rev Neurosci 46, 301–320.

25. Lakshminarasimhan KJ, Avila E, Pitkow X & Angelaki DE (2023). Dynamical latent state computation in the male macaque posterior parietal cortex. Nat Commun 14, 1–20.

26. Lakshminarasimhan KJ, Petsalis M, Park H, DeAngelis GC, Pitkow X & Angelaki DE (2018). A dynamic Bayesian observer model reveals origins of bias in visual path integration. Neuron 99, 194–206.

27. Lappe M, Jenkin M & Harris LR (2007). Travel distance estimation from visual motion by leaky path integration. Exp Brain Res 180, 35–48.

28. Laurens J & Angelaki DE (2017). A unified internal model theory to resolve the paradox of active versus passive self-motion sensation. Elife 6, e28074.

29. Lawrence MA (2016). ez: Easy analysis and visualization of factorial experiments. Available at: https://cran.r-project.org/package=ez.

30. Loomis JM, Klatzky RL, Golledge RG, Cicinelli JG, Pellegrino JW & Fry PA (1993). Nonvisual navigation by blind and sighted: Assessment of path integration ability. J Exp Psychol Gen 122, 73–91.

31. Medendorp WP (2011). Spatial constancy mechanisms in motor control. Philos Trans R Soc B Biol Sci 366, 476–491.

32. Medendorp WP, Alberts BBGT, Verhagen WIM, Koppen M & Selen LPJ (2018). Psychophysical evaluation of sensory reweighting in bilateral vestibulopathy. Front Neurol 9, 377.

33. Medendorp WP & Selen LJP (2017). Vestibular contributions to high-level sensorimotor functions. Neuropsychologia 105, 144–152.

34. Nash CJ & Cole DJ (2020). Identification and validation of a driver steering control model incorporating human sensory dynamics. Veh Syst Dyn 58, 495–517.

35. Nash CJ, Cole DJ & Bigler RS (2016). A review of human sensory dynamics for application to models of driver steering and speed control. Biol Cybern 110, 91–116.

36. Page WK & Duffy CJ (2008). Cortical neuronal responses to optic flow are shaped by visual strategies for steering. Cereb Cortex 18, 727–739.

37. Prsa M, Jimenez-Rezende D & Blanke O (2015). Inference of perceptual priors from path dynamics of passive self-motion. J Neurophysiol 113, 1400–1413.

38. R Core Team (2017). R: A language and environment for statistical computing. Available at: https://www.r-project.org/.

39. Rineau AL, Bringoux L, Sarrazin JC & Berberian B (2023). Being active over one’s own motion: Considering predictive mechanisms in self-motion perception. Neurosci Biobehav Rev 146, 105051.

40. Roy JE & Cullen KE (2001). Selective processing of vestibular reafference during self-generated head motion. J Neurosci 21, 2131–2142.

41. Sanders MC, Chang NYN, Hiss MM, Uchanski RM & Hullar TE (2011). Temporal binding of auditory and rotational stimuli. Exp Brain Res 210, 539–547.

42. Stavropoulos A, Lakshminarasimhan KJ, Laurens J, Pitkow X & Angelaki D (2022). Influence of sensory modality and control dynamics on human path integration. Elife; DOI: 10.7554/eLife.63405.

43. Wei K & Körding K (2010). Uncertainty of feedback and state estimation determines the speed of motor adaptation. Front Comput Neurosci 4, 11.

44. Worchel P (1952). The role of the vestibular organs in space orientation. J Exp Psychol 44, 4–10.

45. Zhou L & Gu Y (2023). Cortical mechanisms of multisensory linear self-motion perception. Neurosci Bull 39, 125–137.

